# Transposable elements activity reveals punctuated patterns of speciation in mammals

**DOI:** 10.1101/082248

**Authors:** Marco Ricci, Valentina Peona, Etienne Guichard, Cristian Taccioli, Alessio Boattini

## Abstract

Transposable elements (TEs) play an essential role in shaping eukaryotic genomes and generating variability. Our “Cold Genome” hypothesis postulates that speciation and TEs activity are strongly related in mammals. In order to test this hypothesis, we created two new parameters: the Density of Insertion (DI) and the Relative Rate of Speciation (RRS). The DI is the ratio between the number of TE insertions in a genome and its size, whereas the RRS is a conditional parameter designed to identify potential speciation bursts. Thus, by analyzing TEs insertions in mammals, we defined the genomes as “hot” (low DI) and “cold” (high DI). Then, comparing TEs activity among 16 intra-order pairs of mammalian species, 4 superorders of Eutheria and 29 taxonomical families of the whole Mammalia class, we showed that taxa with positive RRS correlate with “hot” genomes, whereas taxa with negative RRS correlate with “cold” genomes. In addition, our study supports the “Punctuated Equilibria” theory in mammals for both adaptive radiation and stasis.

## MAIN TEXT

Phyletic Gradualism (PG; Charlesworth et al. 1982) and Punctuated Equilibria (PE; Eldredge and Gould 1972) are the most important evolutionary theories for explaining speciation dynamics. According to PG, species continuously accumulate mutations that would eventually lead to differentiation and speciation. Therefore, older taxonomical groups had more time to accumulate biodiversity, leading to an overabundance of species in compari-son to younger groups (McPeek and Brown 2007). To test the PG hypothesis in Mammalia we calculated the correlation and performing linear regression models between clade age and species richness for 152 mammalian families (**Table S1**). These analyses clearly revealed (**Figure S1**) that there is no significant association between the two variables. Since the PG model does not seem to describe mammalian evolution accurately (**Figure S1**), we hypothesized that the PE theory (already suggested in Mattila and Bokma 2008) and the genomic impact of Transposable Elements (TEs) might better explain their evolutionary dynamics.

TEs are linked to essential cellular activities such as telomere maintenance (Farkash and Prak 2006) rewiring of transcriptional networks (Kunarso et al. 2010), regulation of gene expression (Choung et al. 2016), ectopic recombination and chromosomal rearrangements (Fedoroff 2012) among others. Furthermore, they are key contributors to evolution (Biemont 2010, Oliver et al. 2013, Kapusta et al. 2017) and play a fundamental role in biological processes of utmost importance, like the insurgence of the V(D)J system (Kapitonov and Jurka 2005, Koonin and Krupovic 2014), placental development (Lynch et al. 2011), embryogenesis (Gerdes et al. 2016, Friedli and Trono 2015) and neurogenesis (Richardson et al. 2014). Given their huge impact on shaping genomes, TEs are also thought to contribute to the formation of reproductive barriers facilitating speciation as recently proposed by the Epi-Transposon (Zeh et al. 2009), CArrier SubPopulation (Jurka et al. 2011) and TE-Thrust (Oliver and Greene 2012) hypotheses.

Living organisms owe their ability to diversify to their genomic plasticity and TEs activity and their mutagenic potential can substantially contribute to it (Carmi et al. 2011, Belyayev 2014, Elbarbary et al. 2016, Huff et al. 2016). Hence, we expect that a positive relationship exists between TEs activity and the extant biodiversity. Observing mammalian phylogeny, we can notice that the order Monotremata is the most ancient and the poorest in living species (Figure 1A); accordingly, the platypus genome, belonging to this taxon, harbors the lowest number of recently mobilized TEs (Jurka et al. 2011). Thus is it possible that taxa with low rates of speciation are associated to genomes with mostly inactive TEs?

**Figure 1.**
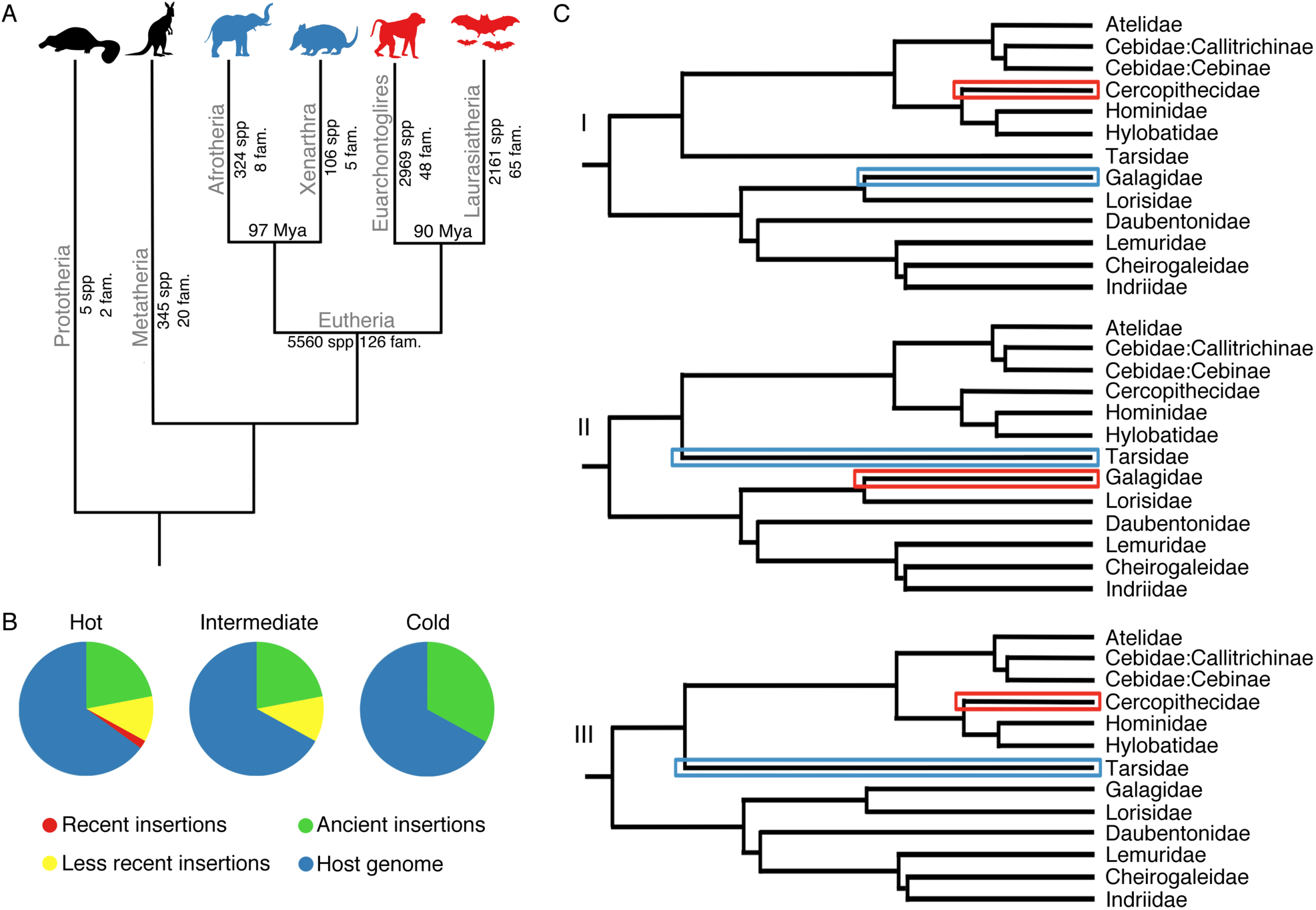
A) Tree of Mammals. Species abundance and phylogenetic relationships of the main mammalian clades. Putatively “hot” superorders of Eutheria (RRS (+)) are shown in red; putatively “cold” superorders (RRS (-)) are shown in blue. Animal icons made by Freepick from www.flaticon.com B) Modelization of the Cold Genome hypothesis. “Hot” genomes contain a fraction of active, recently mobilized TEs (diverging less than 1% from their consensus sequence). “Intermediate” genomes contain a fraction of less recently mobilized TEs (diverging less than 5% from their consensus sequence). “Cold” genomes show ancient insertions with very low or absent activity (like the platypus genome). C) Exemplified use of the Relative Rate of Speciation (RRS) within the order Primates. I Galagidae, when compared to Cercopithecidae, are older and poorer in species, thus Galagidae: RRS (-), Cercopithecidae: RRS (+). II Galagidae, when compared to Tarsidae, are younger and richer in species, thus Galagidae: RRS (+) and Tarsidae: RRS (-). III Cercopithecidae when compared to Tarsidae, are younger and richer in species, thus Cercopithecidae: RRS (+) and Tarsidae: RRS (-). (Color code: RRS (+): red; RRS (-): blue).

Starting from the observations of this specific case, we introduced a general evolutionary model that we call the “Cold Genome” hypothesis. According to it, genomes with highly active TEs (“hot” genomes) belong to taxa with high rates of speciation. The decreasing of TEs activity (intermediate genomes) could lead to “cold” genomes, in which TEs families are almost inactive, therefore to taxa with low rates of speciation (Figure 1B-C). According to our model, the alternations in TEs activity bursts and stasis would be linked to the alternations of bursts and stasis of speciation, giving us a genomic explanation of the PE theory.

To evaluate TEs activity in mammalian genomes, we started from the fundamental study by Jurka et al. 2011 (**Table S2**) that provides the number of insertions and the number of TE families (NF) diverging less than 1% and less than 5% (1%NF, 5%NF) from their consensus (namely reference) sequences. The divergence from the consensus, on a large scale, is a proxy of insertions’ age (Jurka et al. 2011). Thus, insertions diverging less than 1% are more recent, while those diverging 1-5% are older. Subsequently, we created a new parameter called Density of Insertion (DI), which is the ratio between the number of TE insertions in a genome and its size. We calculated the DI at both divergence thresholds (1%DI and 5%DI). As for mammalian speciation patterns, we also calculated the Rate of Speciation (RS) as the ratio between the number of extant species and the crown age (CA) of a taxon (see Materials and Methods). We finally tested the hypothesis that TEs activity is related with speciation patterns by measuring the association between NF, DI and RS (Figure 2) with Spearman correlation and linear regression models (**Table S3-S4**). Notably, all the parameters showed significant correlation with RS in the whole Mammalia class (**Table S3-S4**). In particular, linear models (**Table S4**) showed positive regression coefficients and significant P-values. Therefore, these results suggest a general association between TEs activity and speciation events. It is tempting to speculate that the repertoire of TEs present in a given genome might directly influence its capability to diversify.

**Figure 2.**
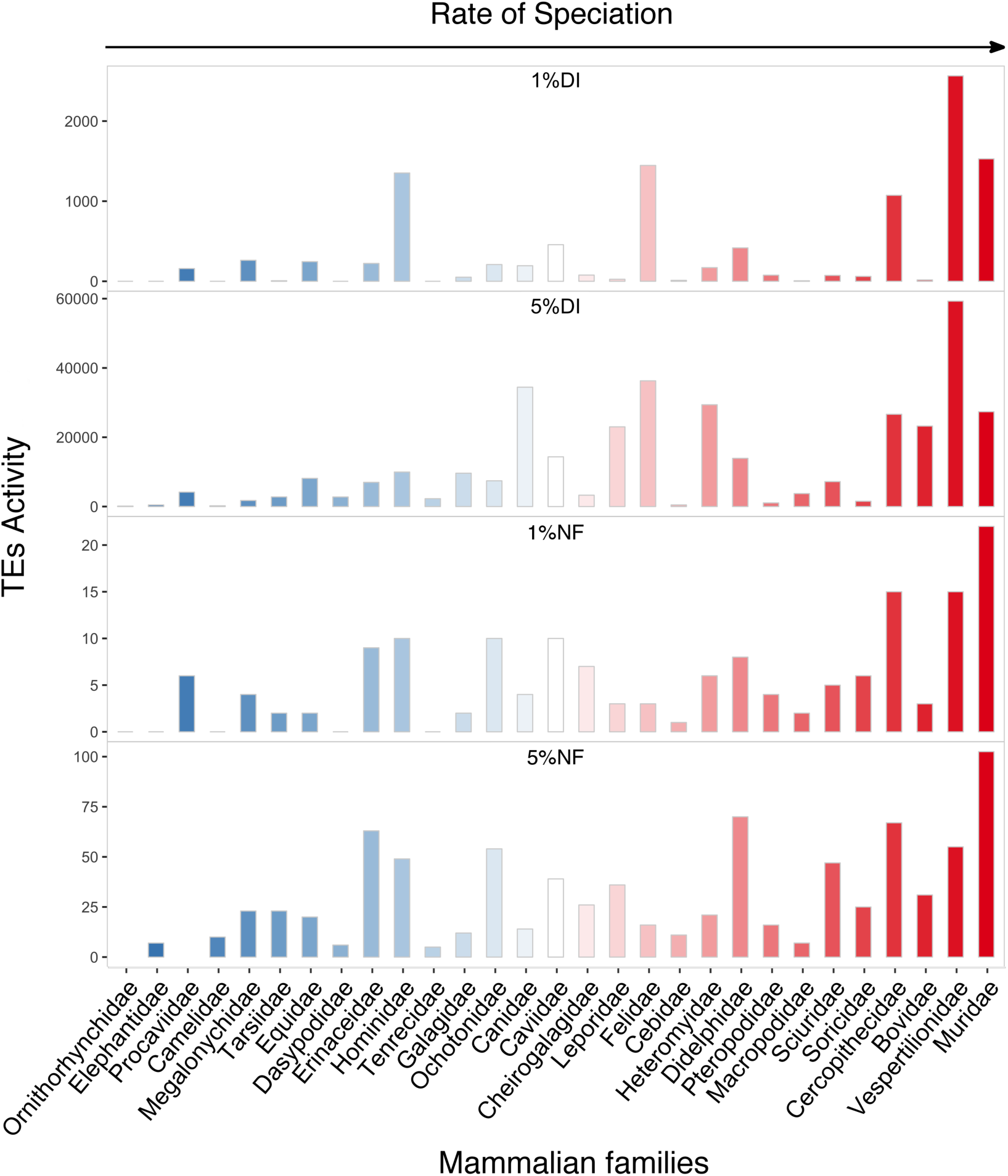
Relationship between the Rate of Speciation (RS) and TEs activity estimated according to the four considered parameters (1%DI, 5%DI, 1%NF, 5%NF)in the 29 mammalian families of Eutheria. The families are arranged in increasing order of RS. (See also **Table S2**, **Table S3** and **Table S4**).

On the other hand, adaptive radiation events can only be defined comparing different taxa and to identify them we designed a new parameter called Relative Rate of Speciation (RRS) (Figure 1C). RRS is a conditional parameter that compares a pair of taxa at same hierarchical level (e.g. two families within the same order). Briefly, if one taxon of a given pair at the same time shows 1) a higher number of species and 2) a lower age compared to its paired taxon, then its RRS is positive (+) and putatively experienced a (relatively) recent speciation burst. Consequently, the other taxon has a negative RRS (-) and is experiencing a more static phase (Figure 1C). If only one of the two conditions is met and there is no evidence of adaptive radiation/stasis for neither of the two taxa (RRS = NA; **Supplementary Text 1**). RRS can be applied at any taxonomical level on any monophyletic clade. In order to minimize potential stochastic noise (e.g. differential TEs activity and/or species richness), in this work we applied the RRS to mammalian families that belong to the same order and to mammalian superorders belonging to the same subclass.

According to our Cold Genome hypothesis, we expect that genomes with higher TEs activity (“hot”) should correspond to RRS (+) taxa, while RRS (-) taxa should have lower TEs activity (“cold” genomes). At the lower taxonomical level herein considered (families within orders), we used 16 mammalian species (encompassing six orders) arranged in 16 pairs (**Table S5**). For each genome, we calculated the four parameters described above (1%DI, 5%DI, 1%NF, 5%NF; **Table S2**). We tested the association between putative “hot”/“cold” genomes (defined via RRS) and TEs activity (DI and NF), with the paired Wilcoxon Signed Rank Test. All tests, excluding 5%DI, were significant, the one with higher confidence being 1%DI (Figure 3A, **Table S6-S7**). Using 1%DI, 14 out of 16 pairs matched the RRS results (**Table S6, Supplementary Text 2**). Furthermore 11 pairs showed a difference in DI of at least one order of magnitude, up to almost 180-fold higher (*Macaca mulatta* vs *Tarsius syrichta*). Accordingly, our results clearly suggest that in Mammals recent TEs activity is associated with recent adaptive radiation. Therefore, the relative level of TEs activity between two taxa is highly related to their relative ability to differentiate and speciate and the activity of TEs does not vary randomly within the mammalian phylogeny. In addition, 1%DI seems to be a more sensible parameter than NF for measuring recent TEs activity (**Supplementary Text 2**).

**Figure 3.**
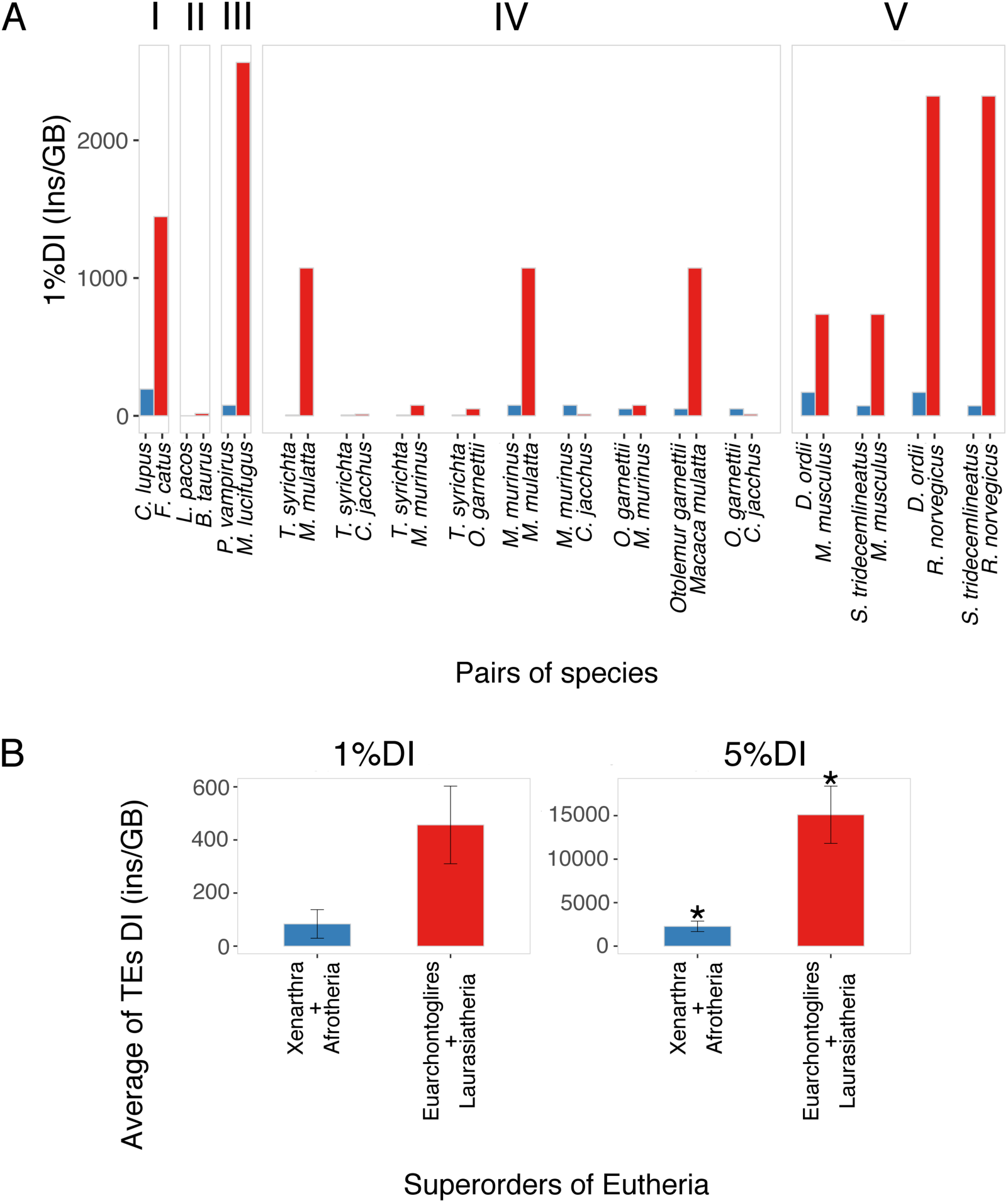
**A)**1%DI values in the 16 pairs of mammalian species which exhibit evidence of adaptive radiation/stasis. Blue bars: RRS (-) (putative “cold” genomes); red bars: RRS (+) (putative “hot” genomes). **I** Carnivora, **II** Cetartiodactyla, **III** Chiroptera, **IV** Primates, **V** Rodentia. **B)** Comparison of the 1%DI and 5%DI in the 4 superorders of Eutheria. Blue bars: RRS (-) (putative “cold” genomes); red bars: RRS (+) (putative “hot” genomes). * P-value < 0.05.

Next, we tested our hypothesis at a higher taxonomic level considering the four Eutheria superorders of Afrotheria (A), Euarchontoglires (E), Laurasiatheria (L) and Xenarthra (X) (Figure 3B, **Table S8**). According to RRS results, E and L showed RRS (+), thus putatively they are “hot” taxa, while A and X showed RRS (-), thus putatively they are “cold” taxa (Figure 1A). Accordingly, after averaging their respective DIs, we merged the putatively “hot” superorders (E and L, 22 species) and the putatively “cold” superorders (A and X, 5 species) and tested their association with DI as above (**Supplementary Text 3**). For both 1%DI and 5%DI, “cold” superorders show an averaged DI more than three-fold lower than “hot” superorders. Differently from what observed at the lower taxonomical level, 5%DI yielded a significant difference between the two groups, while the test with 1%DI is nonsignificant (Figure 3B).

This discrepancy between the lower and higher taxonomical levels may be interpreted from an evolutionary point of view. The 5%DI is the worst predictor of TEs activity among the four considered parameters (1%DI, 5%DI, 1%NF, 5%NF) when studying recent speciation (Figure 3A, **Table S7**). On the contrary, it is the best one when considering older macroevolutionary events, such as the differentiation of the four Eutheria superorders (Figure 3B, **Table S9**). Hence, the divergence of the elements from their consensus reflects, in average, their age and related adaptive radiation events.

In conclusion, our results suggest that TEs activity may influence speciation patterns in Mammals. In fact, a high differentiation rate in a taxon is strongly associated with an increased molecular activity of the TEs (see also Feiner 2016). Moreover, TEs seem to be important for adaptive radiation.

In evolutionary time-scales, we hypothesize that their activity is modulated, producing alternations of insertional bursts and silencings (Muñoz-Lopez et al. 2011). Accordingly, recent studies show that young LINE-1 elements are mostly repressed via methylation while old TEs are regulated by the KRAB/KAP1 system (Castro-Diaz et al. 2015).

While silencing mechanisms progressively inhibit TEs activity (state of “cold” genome), their lack of contribution to molecular differentiation might lead to the relatively static phase postulated by the PE theory.

Thus, the “Cold Genome” hypothesis seems to support the PE theory in both the punctuated differentiation bursts and stasis periods. Furthermore, we showed that TE insertions and their approximate occurrence times are in accordance with clade differentiation: older TE bursts are associated to older adaptive radiation events (origin of mammalian superorders), whereas novel TE bursts correlate to newer evolutionary phenomena (origin of mammalian families). Whether TEs mobilization and accumulation of new insertions is the cause or the effect of adaptive radiation/speciation remains open for debate. However, the results presented here and the intrinsic characteristics of the mobilome activity suggest that TEs might play an important role in molecular differentiation of living organisms. Further studies associating TEs and speciation can shed new light on our understanding of evolutionary processes.

## MATERIALS AND METHODS

The number of species for all 152 mammalian families listed in the last mammalian phylogeny (Meredith et al.2011) was retrieved from Catalogue of Life (http://www.catalogueofli-fe.com); their crown age (CA) was estimated from their timed phylogenetic tree (Meredith et al. 2011). Data about TE families and TE insertions in the genomes of the considered species was retrieved from the work of Jurka et al. 2011.

DI is calculated according to the formula: DI=NI/GS, where NI is the total Number of Insertions (of elements contained in the 1% or 5% datasets) and GS is the Genome Size in Gigabases.

RS is calculated with the formula: RS=NS/CA, where NS is the total number of species for the analyzed taxonomical family.

The RRS attribution can be represented by the logical formulae:

RRS1(+): NS1 > NS2 ^ CA1 < CA2
RRS1(-): NS1 < NS2 ^ CA1 > CA2

If one of these conditions is false, there is no evidence of adaptive radiation events between the considered taxa therefore RRS = 0.

RRS was applied on families belonging to the same order (**Table S5, Supplementary Text 2**) and to the four superorders of Eutheria (**Table S8, Supplementary Text 3**).

We tested the correlation between putative hot/cold genomes and RRS (+/-) using the Wilcoxon Signed Rank Test either for both families and superorders. All statistical analyses and graphs were performed/produced with the R software (R Core Team 2016).

